# Graphicacy across age, education, and culture: a new tool to assess intuitive graphics skills

**DOI:** 10.1101/2022.10.24.513063

**Authors:** Lorenzo Ciccione, Mathias Sablé-Meyer, Esther Boissin, Mathilde Josserand, Cassandra Potier-Watkins, Serge Caparos, Stanislas Dehaene

**Affiliations:** Cognitive Neuroimaging Unit, CEA, INSERM, Université Paris-Saclay, NeuroSpin center, 91191 Gif/Yvette, France; Collège de France, Université Paris Sciences Lettres (PSL), 75005 Paris, France; LaPsyDÉ, CNRS, Université Paris Cité, F-75005 Paris, France; Laboratoire Dynamique Du Langage UMR 5596, Université Lumière Lyon 2, 69363 Lyon, France; Department of Psychology, DysCo lab, Université Paris 8, 93526 Saint-Denis, France; Human Sciences Section, Institut Universitaire de France, 75005 Paris, France

## Abstract

Data plots are widely used in science, journalism and politics, since they efficiently allow to depict a large amount of information. Graphicacy, the ability to understand graphs, thus became a fundamental cultural skill. Here, we introduce a new measure of graphicacy that assesses the ability to detect a trend in noisy scatterplots (“does this graph go up or down?”). In 3943 educated participants, responses vary as a sigmoid function of the *t*-value that a statistician would compute to detect a significant trend. We find a minimum level of core graphicacy even in unschooled participants living in remote Namibian villages (N=87) and 6-year-old 1^st^-graders who never read a graph (N=27). However, the sigmoid slope (the “graphicacy index”) varies across participants, increases with education, and tightly correlates with statistical knowledge, showing that experience contributes to refining graphical intuitions. Our tool is publicly available online and allows to quickly evaluate intuitive graphics skills.

**STATEMENT OF RELEVANCE:** The rising cost of gas, the number of Covid deaths, the evolution of temperatures during the summer months: we often face graphs depicting these phenomena. The scientific literature has shown that human adults can intuit, within milliseconds, the statistical trend of these graphs. However, we do not know if these intuitions generalized to unschooled people and, most importantly, how to measure their variations in the population. In this study we show that intuitive graphics skills are present even in 6-year-old children who never saw a graph and in the Himba of Namibia, an indigenous people with no access to formal schooling. Furthermore, we developed a quantitative assessment of such intuitive graphics skills (which we called the “graphicacy index”), that everyone can easily obtain for free, through a short (10 minutes) online test: https://neurospin-data.cea.fr/exp/lorenzo-ciccione/graphicacy-index/. In summary, our study provides the first attempt to formally quantify human intuitions of statistical graphs.

## INTRODUCTION

Humans often exhibit a surprising intuitive grasp of the core concepts of mathematics, physics or statistics. These intuitive abilities, which emerge in the absence of formal education, are likely to rely on a system of core implicit knowledge about the fundamental properties of the environment in which humans evolved (Spelke & Kinzler, 2007). A solid body of research shows, for example, that humans can accurately and quickly grasp the approximate numerosity of sets of objects (Dehaene, 2011) and perform approximate calculations even in the absence of formal mathematical education (Pica et al., 2004). Euclidean and non-Euclidian geometrical intuitions of space are present in remote Amazon populations without access to formal education (Dehaene et al., 2006; Izard et al., 2011). In concrete settings, humans also excel in intuitive physics: their misconceptions about moving objects’ behavior (McCloskey, 1983) disappear when questions are framed in familiar and real-life contexts (Kubricht et al., 2017). Humans are also remarkably good at performing intuitive statistical estimations in a variety of tasks (Nisbett & Krantz, 1983) and they are endowed with these abilities from early on in their development (Xu & Garcia, 2008). Indeed, many quantitative assessments of intuitive mathematics and physics exist, and they often predict the subsequent development of higher-level cognitive abilities (Baron-Cohen et al., 2001; Halberda et al., 2008; Perez & Feigenson, 2021; Piazza et al., 2010; Riener et al., 2005), suggesting a strong link between basic intuitions and the formal mastery of complex concepts.

Whether these core intuitions extend to graphical representations is still an open question. In fact, despite the widespread use of charts and plots in our everyday life, the human competence for graph perception (graphicacy) remains vastly understudied, and no quantitative assessment of intuitive graphics skills has been proposed. Here, we show how the recent development of graph-based psychophysics tasks offers a tool to quantify intuitive graphics skills. We previously showed that, when facing a graph such as a noisy scatterplot (figure 1A), human adults can detect whether the curve is increasing or decreasing, regardless of the number of dots, noise level or slope of the graph. Their performance is predicted by the *t*-value that a statistician would calculate to determine the significance of the trend in the data (Ciccione & Dehaene, 2021). In other words, the percentage of “increasing” responses is a sigmoid function of the *t*-value of the scatterplot. Here, we show that those findings are highly reliable across a variety of populations and can be used to assess graphicacy. As in any two-alternative forced-choice task, the slope of the psychometric function (figure 1B) provides a measure of a participant’s sensitivity to detect variations in the stimulus: the steeper the function, the higher the participant’s precision. We demonstrate that this simple psychophysical measure provides a simple quantitative assessment of how good a single individual is in graph perception.

**Figure 1.**
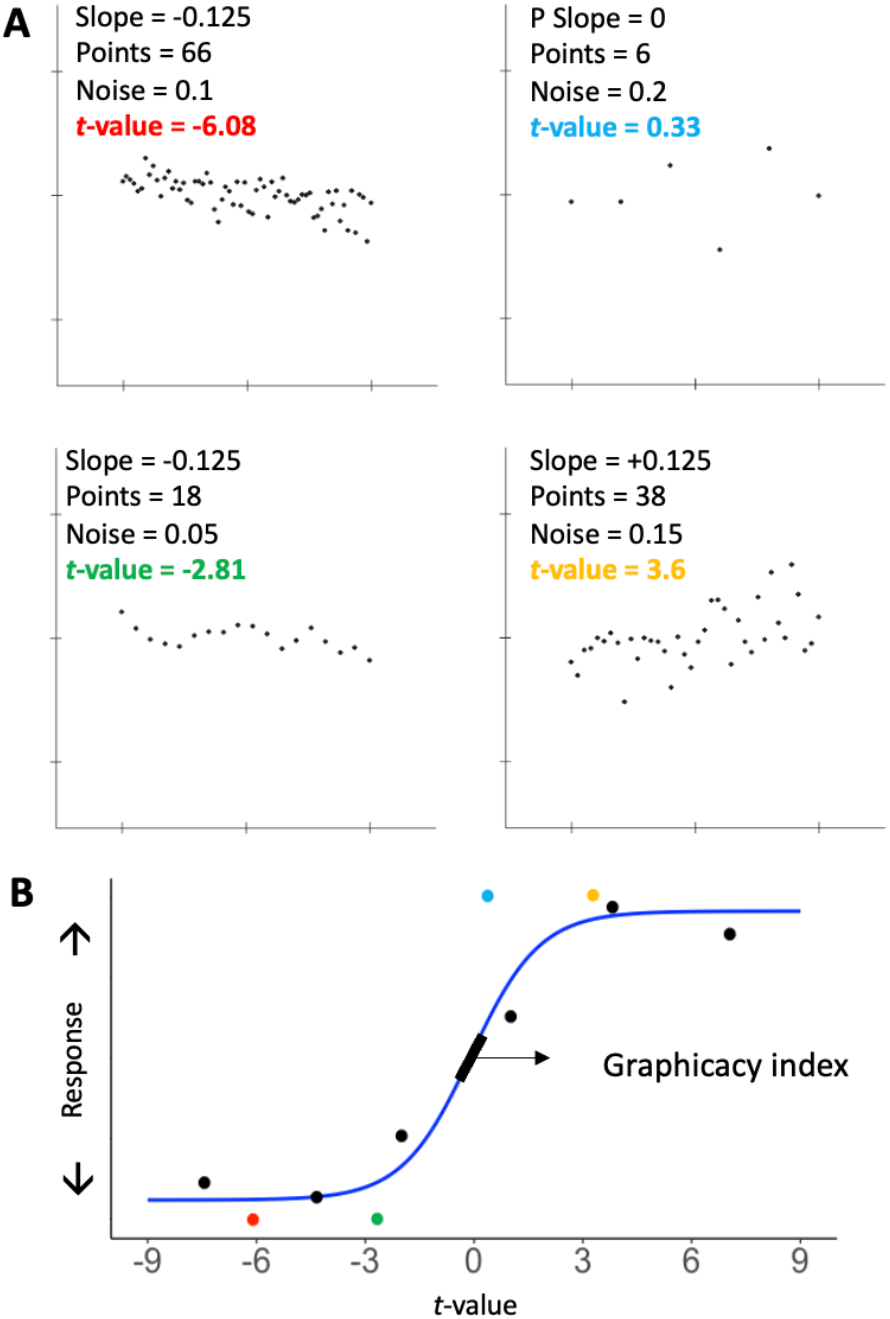
A simple test of graphicacy. **A:** four examples of stimuli shown to one participant in the trend judgment task, where the participant was asked if the graph was increasing or decreasing. The actual stimuli were white dots on a black background. Each scatterplot was created according to the combinations of different parameters: slope of the generative function, number of points, and noise level. Each thus had a certain t-value, corresponding to the *t* statistic used to calculate the significance of the trend in the dataset. **B:** Responses given by a representative participant are plotted as a function of the *t*-value of each scatterplot. Color dots show the data for the four example trials in panel **A**. For visualization purposes, the black dots show averages over bins of t-values. We fitted the data with a psychometric function (blue curve). The slope of the sigmoid, indicated by a black bar, evaluates the subject’s sensitivity in the trend judgment task, and was used as a “graphicacy index”, a proxy of the participant’s intuitive graphics skills.

In the present work, we investigated the emergence and distribution of trend judgment skills across people of different ages, education levels and cultures. First, we tested intuitive graphics in a large-scale online sample of educated adults from all over the world, from which we obtained information about several demographical aspects, together with self-reports of mathematical and statistical understanding. Testing intuitive graphics on such a large and diverse population offered insights into the predictors of these skills; also, it provided a large-scale replication of psychophysical results previously obtained with a reduced sample in a controlled laboratory environment (Ciccione & Dehaene, 2021), thus contributing to the growing but still scarce body of research on psychophysical measurements outside the lab (e.g., Ciccione et al., 2022; de Leeuw & Motz, 2016; Halberda et al., 2012; Semmelmann & Weigelt, 2017).

Second, we explored whether graphicacy emerges as a result of graph exposure or whether some premises of graphicacy are available even in the absence of formal education. To investigate this, we tested Himba participants, a Namibian people with no or little formal education, who are not exposed to any form of graphical representations. This sample of participants allowed us to test for the generalizability of such skills in non-western and unindustrialized societies, as previously been done for other intuitive skills (Spelke & Kinzler, 2007), including the perception of number (Pica et al., 2004) and geometry (Dehaene et al., 2006; Izard et al., 2011; Sablé-Meyer et al., 2021).

In addition, we tested graphicacy in French 6-year-old 1^st^-graders who never encountered any graphical representation in their school curriculum. We thus asked whether the ability to compute intuitive visual statistics from graphs arises early on in development, as should be the case if it relies on a core skill of human cognition, similar to number sense or shape perception. The cultural recycling hypothesis (Dehaene & Cohen, 2007) postulates that the evolutionary old cognitive functions of numerosity and shape perception serve as a foundation for the corresponding culturally learned skills (respectively, arithmetic and reading). Similarly, humans’ ability to read and interpret complex graphs might be based on fundamental cognitive functions available early on in development, irrespective of formal education, such as the recognition of the orientation and medial axis of objects (their “shape skeleton”; Feldman & Singh, 2006; Firestone & Scholl, 2014; Lowet et al., 2018). Previous results point indeed to that direction: simple trend judgments performed on noisy scatterplots rely on an estimation of their principal axis (Ciccione & Dehaene, 2021). Showing that graphical intuitions are available to 1^st^-graders, prior to any formal education in mathematical graphs, would constitute further evidence of their core nature.

## METHODS

### Experimental procedure and participants

#### Online participants

The online test was advertised and shared on social networks, mainly through Twitter. It could be performed either on computers or on tactile devices. Before taking part in the experiment, participants had to read and accept a written consent and to declare to be at least 18 years old, in compliance with the local Ethical Committee that approved our research (under the reference CER-Paris-Saclay-2019-061). Data collection for the purpose of the study started on January 15^th^, 2021 and ended on March 15^th^, 2021, as planned ahead of the experiment. The link to the test was still running after that date, but the data were not included in the current work.

Before taking the test, all participants answered a demographic questionnaire consisting in a series of single-answer questions about: country of origin, age, gender, number of previous participations in the task (if any) and the highest level of education attained. If participants declared to have completed a university degree, they were asked to choose the closest field of the degree within a list, and their average grade in mathematics during their school and university years. Using a Likert scale (ranging from 1 to 10, with intermediate numbers not shown), all participants had to rate their subjective self-evaluation in the following domains: familiarity with graphs, ability to read scatterplots, knowledge of statistics, current skills in mathematics, and current skills in their first language in terms of spelling, grammar and communication. Once the demographic questionnaire was completed, participants started the experiment. Smartphone users were asked to rotate their phone horizontally: otherwise, the task would not start; accidentally orientating the phone vertically during the task lead to a pause in the experiment. The instructions and the questionnaire were available in six languages: English, French, Italian, Spanish, Portuguese and Chinese. 3943 subjects participated and completed the online experiment (the ones that did not complete the task were not included in the data analysis). 2409 of them declared being women, 1294 men, 82 non-binary, 20 “other” than the previous ones, and 138 preferred not to answer. The average age was 28.8 ±9.6 years.

*Himba*. 87 Himba participants (39 women and 48 men) were recruited in small villages in the Kunene region, Northern Namibia. Most Himba do not know their age. Participants’ age, 21.1 ± 9.4 years, was evaluated by local research assistants who were bilingual Namibians (in Otjiherero and English). Those assistants also instructed each participant, in their native language (Otjiherero), about how to perform the task on a tablet. Before the experiment, each participant was provided with four examples of stimuli and the expected correct answers. Each participant indicated whether they had received any type of formal schooling. Rudimentary mobile schools (using black board and chalk) exist in the Kunene region, and only 12 participants declared having received at least one year of such form of schooling.

#### Children

27 French 1^st^ graders (6 ± 0.6 years; 13 girls) took part in the experiment (approved by the local ethical committee under the reference CER-Paris-Saclay-2021-046) and completed the experimental tasks. Each child was accompanied by an experimenter to a silent room and invited to sit on a chair facing a table. Before starting the actual experiment, they performed three short behavioral tests: a one-minute reading task consisting in a series of French words of increasing difficulty; a one-minute counting task on sets of points of increasing numerosity; and a one-minute counting task on those same sets of points, but organized in groups (e.g., 4 groups of 3). The first task provided a number of correctly read items in one minute, which was used as a proxy of reading abilities. The difference in correctly enumerated items between the second and the third task provided an implicit measure of the mastery of arithmetic operations, because grouped items can be enumerated faster if children know how to perform mental arithmetic (“groupitizing”; Ciccione & Dehaene, 2020; Starkey & McCandliss, 2014). The main experimental task was performed on a tablet and, immediately before it, each child was provided with four examples of stimuli and their expected correct answers. The sample size of Himba and children participants was based on a previous study that used an identical trend judgment task (Ciccione & Dehaene, 2021). All data and scripts for the analyses are available on the Open Science Framework at: https://osf.io/cw6t5/.

### Experimental task

Each trial consisted in the rapid presentation (100 ms) of a scatterplot (figure 1A). Participants performed a trend judgment task: they had to judge, as fast and accurately as possible, whether the scatterplot was increasing or decreasing by pressing one of two separate keys on their computer keyboard or, if they played on a smartphone/tablet, by touching an upwards or a downwards arrow. For the online experiment, the response configuration of the keys and the arrows was randomly determined at the beginning of the experiment for each subject, in order to control for possible preferential response sides; also, each correct response was rewarded with a certain amount of points, inversely proportional to the response time. Such gamification incited participants to be both accurate and fast. To maintain a high level of attention in the task, consecutive correct responses were rewarded with increasingly higher points. Also, a pleasant sound followed each correct trial and an unpleasant sound followed each incorrect trial. For children and Himba participants, a smiling green face or a red unsmiling face was displayed instead of the numerical score. A fixation cross was presented for 1000 ms before the following trial appeared. The experimental session lasted around 6 minutes. Online and Himba participants had the opportunity to start another run or to stop. Online participants could also check their percentage of correct responses and their ranking relative to all previous participants. For data analysis, we rejected any answer that was given after more than 5 seconds from stimulus onset (0.75% of trials for online participants; 9.39% for children; 0.91% for the Himba).

### Stimuli

The stimulus generation algorithm was identical to the one used in a previous laboratory version of the task (Ciccione & Dehaene, 2021). Each scatterplot was the graphical representation of a dataset randomly generated from a linear equation of the form y_i_ = α x_i_ + ε_i_, where α is the prescribed slope and the ε_i_ are random numbers drawn from a normal distribution centered on zero and with standard deviation +. A total of 112 scatterplots were presented to participants, which were the result of the combinations of 3 orthogonal factors: 7 prescribed slopes (α = −0.1875, −0.125, −0.0625, 0, +0.0625, +0.125 or +0.1875); 4 levels of noise (σ = 0.05, 0.1, 0.15 or 0.2); and 4 numbers of points (n = 6, 18, 38, 66). All coordinates on the x axis were fixed and equally spaced for each level of n. Figure 1A shows four examples of stimuli. For each scatterplot, the *t*-value associated to its Pearson coefficient of correlation was calculated. Figure 1 shows examples of stimuli and responses for one subject. Answers are plotted as a function of the *t*-value of the corresponding trial. For each subject, we fitted a classic psychometric function to the data (shown in blue in figure 1A) and we extracted its slope, which provided a measure of precision at performing the trend judgment task. The first 12 trials for each subject were considered as practice trials and thus excluded from the computation of this index. Also, a minority of subjects that participated in the online experiment had a very large sensitivity index, meaning that their performance was close to perfect (in fact, it was better modelled by a step function rather than by a sigmoid one). To avoid excessive variability, sensitivities higher than 5 (.03% of all participants) were capped at 5.

## RESULTS

### Performance in the trend judgment task is predicted by the *t*-value of the scatterplot

First, we looked at the percentage of “increasing” responses as a function of the prescribed slope (i.e., the steepness of the scatterplot), the noise level and the number of dots (figure 2). We replicated results from previous research conducted on a small sample of subjects in a laboratory context (Ciccione & Dehaene, 2021), finding that the proportion of responses “increasing” was affected by all the above parameters (figures 2A and 2B). In an ANOVA on the proportion of “increasing” responses as a function of the prescribed slope, the noise level and the number of points, all factors had a significant main effect, and the prescribed slope significantly interacted with both the noise and the number of points (all p < .001). These findings confirm what is clearly visible in figures 2A and 2B: the smaller the slope of the graph, the higher the influence of the noise level and the number of points on the trend judgment task. No interaction effect was found between the noise and the number of points (F[8.5, 19667.8] = .81, p = .6), suggesting that the two factors independently affected human trend judgments.

**Figure 2.**
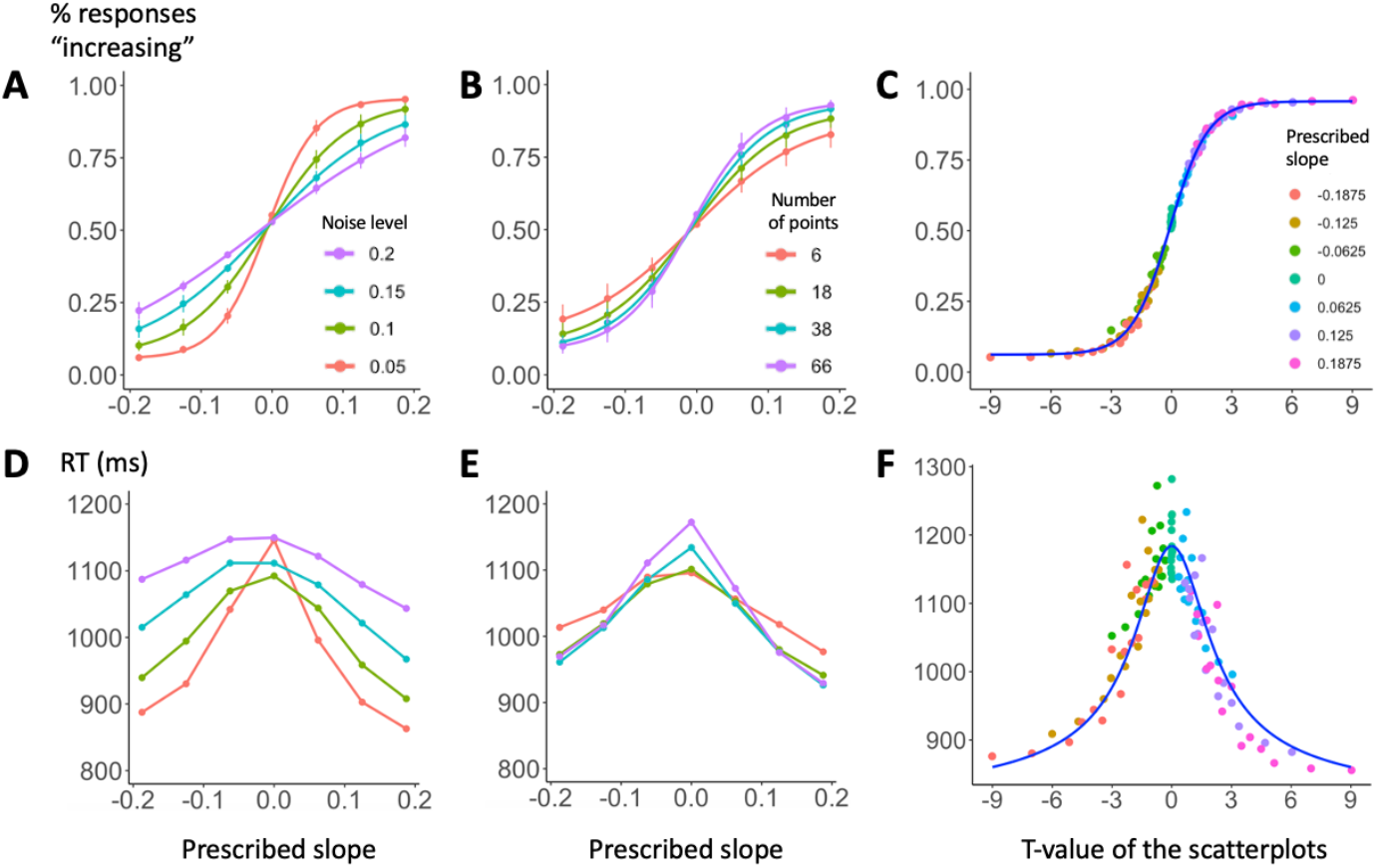
Psychophysics of graph perception (N=3943). The top row shows the percentage of responses “increasing” as a function of prescribed slope and noise level (A), prescribed slope and number of points (B), and *t*-value associated to the Pearson correlation coefficient of the scatterplot (C). Panel C shows that participants’ performance can be subsumed by the t value. D-F, equivalent plots for response time. The blue line in plot F shows the prediction of a simple accumulation of evidence model (Gold and Shadlen, 2002).

All of these effects, however, were subsumed by an effect of the *t*-value associated to the Pearson coefficient of correlation (figure 2C), which varies positively and linearly with the prescribed slope, positively with the number of points (as the square root of n-2) and inversely with the noise level. Accordingly, we computed a multiple logistic regression on “increasing” responses as a function of the *t*-value, the number of points and the noise level (averaged across the 112 combinations of the experimental conditions and across all subjects), and we found that only the *t*-value was a significant predictor of participants’ responses (β_t-value_= .77, p < .0001; β_number of points_= .002, p = .83; β_noise_= −.48, p = .91).

The classical t test formula implies that t grows as the square root of the number of points minus 2. In anticipation of this rule, we used four values for the number of points that were linearly spaced after this square-root transformation (n = 6, 18, 38, 66) and, indeed, we found that performance was roughly equally spaced according to this parameter (figure 2B).

As far as response times for correct answers are concerned (figures 2D and 2E), we submitted them to separate linear regressions as a function of the three main experimental factors (the absolute value of the main slope, the noise level and the number of points) and with the subjects as random factors. We found that, while both the prescribed slope and the noise level significantly predicted the response times (β_slope_ = −946.5, p < .0001; β_noise_ = 920.3, p < .0001), this was not the case for the number of points (β_number of points_ = .06, p > .05), thus suggesting a parallel processing of all items in the set. A simple model of noisy evidence accumulation (Ciccione & Dehaene, 2021; Gold & Shadlen, 2002; see those papers for modeling details) correctly fitted the average response times on all trials (blue line in figure 2F), based solely on the percentage of responses given by the subjects.

### The trend judgement task provides a reliable index of graphicacy and its variations across individuals

We modeled participants’ responses as a sigmoid function of the *t*-value of each stimulus they saw (figure 1B). We postulated that the steepness of this function reflects their intuitive graphics skills. Thus, we called the slope of the sigmoid for a given participant his or her “graphicacy index”. Figure 3A shows the distribution of this index across the large sample we collected online (median value = 1.26). For the vast majority of participants (98.2%), the regression was significant and with a positive index, thus providing a reliable estimate. However, graphicacy varied considerably, with 95% of the distribution falling between 0.20 and 3.18.

**Figure 3.**
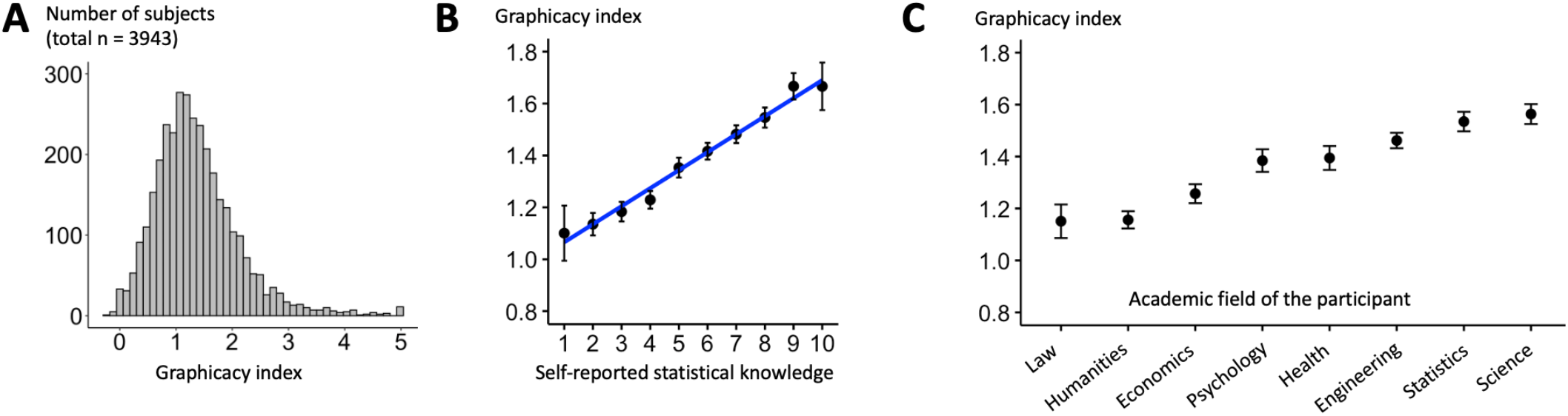
Inter-individual variability in graphicacy. **A:** Distribution of the graphicacy index cross participants. **B:** Graphicacy increases as a function of participants’ self-reported statistical nowledge, collected before the experimental task was introduced. **C:** Graphicacy in participants that obtained at least a bachelor degree varies as a function of the academic field in which they graduated (F(7,2891)= 15.57, p<.001; note that data were ordered according to each group’s mean graphicacy index).

To evaluate the stability of this index during the course of an experimental run, and test for training effects, for the 3419 participants that performed only one experimental run, we computed the index separately on the first 50 trials and on the following 50 trials. Although the increase was significant (Wilcoxon signed rank test, p <.0001), it was small, passing from a median of 1.28 to 1.35. Most crucially, there was a significant correlation between the two values (r(3417) = .38, p < .0001), thus showing a stability of inter-individual variations in graphicacy. To further evaluate whether the graphicacy index remained stable over time, we computed the slope of the orthogonal (Deming) linear regression between those two measurements and we found that the regression slope was close to one (1.02; 95% confidence interval = [.94, 1.1]), thus suggesting that, on average, the index was quite stable and reproducible within an individual.

We also analyzed those subjects that completed more than one block of trials (n = 387). The correlation between their graphicacy index in the first experimental run and in the second one was high (r(385) = .49, p < .0001) and the slope of the orthogonal linear regression between them was again close to one (1.16; 95% confidence interval = [.95, 1.36]).

Overall, these results suggest that our measure of intuitive graphics skills is stable, at least in the absence of extensive training, and can be reasonably estimated in a 6-minute on-line test. It is likely that, should one require a more stable individual measure, a longer testing session would provide an even more reliable graphicacy index.

### Graphicacy correlates with statistical knowledge and academic field

We then tested whether graphicacy correlated with participants’ self-evaluation of their statistical knowledge.

We found a significant correlation (figure 3B; r = .21, df = 3092, p < .0001) between participants’ graphicacy index and their self-reported statistical knowledge. How specific was this correlation? A large majority of subjects (N = 2030) also answered a self-evaluation question on their first language skills, always using a scale from 1 to 10. We performed a multiple linear regression on the graphicacy index as a function of statistical knowledge and language skills, finding that the former was a significant predictor (β = .07, p < .0001) but the latter was not (β = −.006, p = .55). This finding suggests that participants’ graphicacy was not simply predicted by general personal skills, or self-confidence.

Figure 3C shows how the mean graphicacy index varied as a function of the academic field in which graduate participants obtained their title (F(7,2891)=15.57, p<.001): it was considerably higher for graduates in engineering, statistics and science (n = 1576, mean = 1.1) than for graduates in other disciplines (n = 1323, mean = 1.27; t(2894.4) = 8.92, p <.0001). In graduate subjects, the graphicacy index also significantly correlated with their reported average grade in mathematics (r = .24, df = 3028, p < .0001).

Correlations between the graphicacy index and other factors are briefly presented in this appendix.

### Himba performance is predicted by the *t*-value of the scatterplot, independently of age and education level

Figure 4 (left plot) shows that Himba performance in trend judgment was again well predicted by the *t*-value of the scatterplot. We computed a multiple logistic regression of responses “increasing” as a function of the *t*-value, the number of points and the noise level (averaged across the 112 combinations of the experimental conditions and across all subjects), and we found that, again, the *t*-value was the only significant predictor of participants’ responses (β = .27, p < .01), while the noise (β = .37, p = .92) and the number of points (β = .001, p = .86) were not. The same findings held when we separated our data in three separate groups (figure 4, middle plots): teenagers (i.e., participants younger than 18 years old, N=36), unschooled adults (i.e., participants who did not receive any formal education, N=39), and schooled adults (i.e., participants who attended mobile schools during at least one year, N=12). For all these subgroups, responses were entirely accounted for by the *t*-value of the stimulus (all β with p < .01). The median graphicacy index for Himba was of .32, i.e., on the very low end of the distribution of Western educated subjects (see figure 3A).

**Figure 4.**
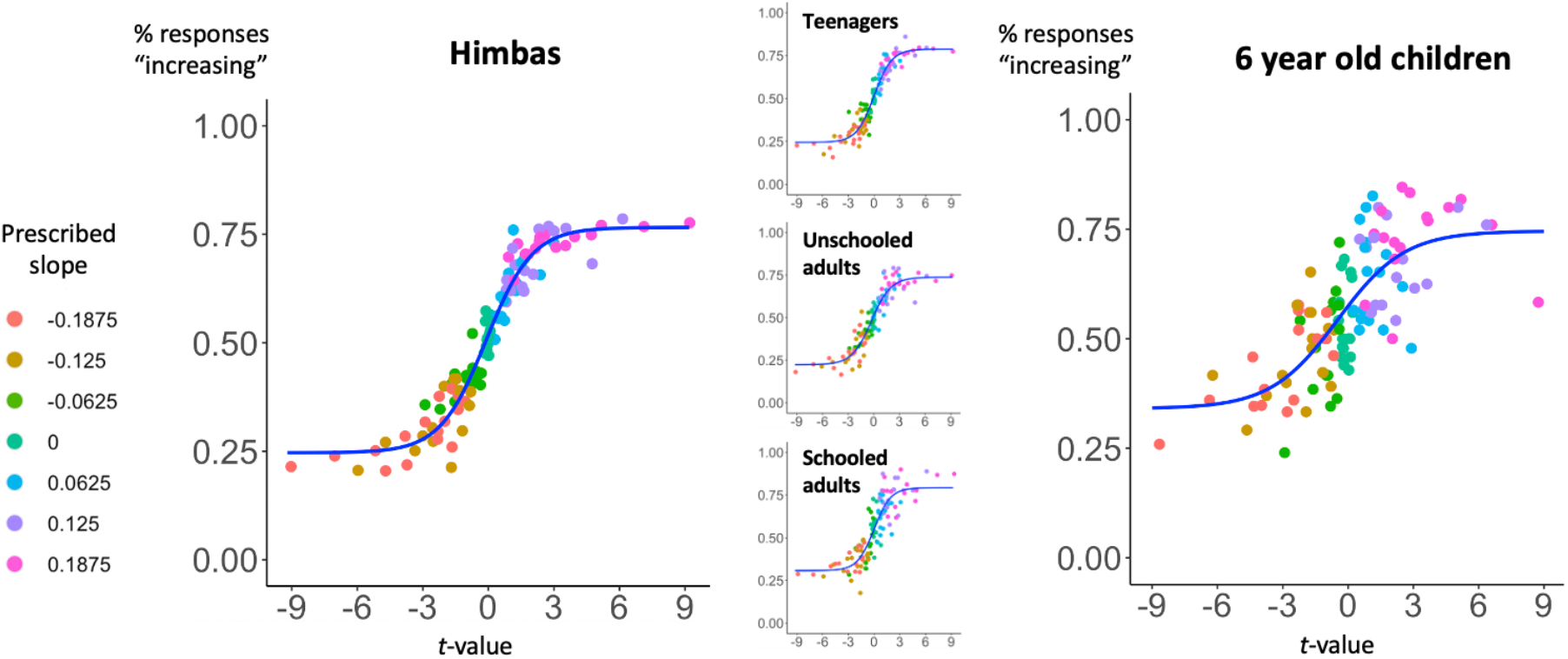
Precursors of graphicacy in the absence of formal education. The percentage of responses “increasing” is well predicted by the *t*-value associated with the graph Pearson correlation coefficient *r*, both in 6-years-old children (right, N=27) and in the Himba participants (left, N=87), an ethnic group from Namibia with reduced access to formal education. Insets show the effect separately as a function of schooling and age for the Himba.

### 6-years-old children’s performance is also predicted by the *t*-value of the scatterplot

The results were also replicated in a group of 27 6-years-old 1^st^-grade children (figure 4, right plot). Although children’s average responses were noisier and never reached perfect performance (as is clear from the boundaries of the sigmoid function in figure 4, right plot), their responses were again significantly predicted by the *t*-value of the scatterplot, which sufficed to account for children’s performance: it was a significant predictor of their responses (β = .17, p < .05), whereas the noise (β = 1.97, p = .58) and the number of points (β = −.0003, p = .98) were not.

For each child, we computed the graphicacy index and correlated it with the two measures described in the methods’ section: groupitizing advantage (an implicit measure of their arithmetic abilities) and reading speed (number of correctly read words in one minute). Both correlations were significant (respectively: r = .51, df = 25, p < .01, and r = .46, df = 25, p < .05). It is worth noting that groupitizing and reading speed were also highly correlated (r = .7, df = 25, p < .0001). The median graphicacy index for children was of .12.

## DISCUSSION

In our study, we measured the human ability to perform a trend judgment task on a noisy graph (i.e., “Does this graph go up or down?”). Analyzing the responses of 3943 participants that performed the task online on computers or tactile devices, we found that their accuracy was affected by all three manipulated factors, namely, the steepness of the graph, its noise level and the number of points. In terms of response times, there was a significant effect of steepness and noise but not of the number of points. This finding suggests that participants treated the scatterplots as an ensemble, without serially processing each item: in fact, if that was the case, we should have observed an increment in response times proportional to the number of points in the dataset. Fast intuitive statistical judgments on graphs with Gaussian noise thus seem to operate similarly to ensemble perception, the human ability to rapidly extract the “average” of visually displayed items, without focusing on each particular element in the set (Cui & Liu, 2021; Whitney & Yamanashi Leib, 2018; but see, in the presence of outliers in the graph: Ciccione et al., 2022).

Crucially, participants’ responses were entirely predicted by the *t*-value associated to the Pearson coefficient of correlation of the graph, showing that humans’ trend judgments approach those of an optimal statistical model (Peterson & Beach, 1967). Our findings thus challenge the view according to which people are naïve, biased or inaccurate intuitive statisticians. While some data show that people do not include sample size in their variability estimations (Kareev et al., 2002), thus suggesting that they wrongly assume that a small sample is always representative of the entire population (Fiedler, 2000; Juslin et al., 2007), our studies demonstrate that, for datasets represented in a bivariate graphical format, people correctly incorporate both variability and sample size in their trend judgments. In other words, at least at a perceptual level, humans are not naïve in their statistical estimates, but seem to take into account all the parameters of the dataset.

It is worth noting, tangentially, that our online study replicated the findings from an identical trend judgment task performed in a laboratory context (Ciccione & Dehaene, 2021). This piece of evidence is important both empirically and methodologically. It confirms that, unlike popular opinion among many scientists, psychophysical studies do not need to be confined to a controlled laboratory environment and can be successfully performed online. This clearly reduces research times and costs, especially when participants, such as in the present online study, were included on a purely voluntary basis and with no other reward than personal enjoyment.

The first drive of our study was to introduce a quantitative measure of intuitive graphics skills, the graphicacy index. We operationalized it as the slope of the psychometric function linking subjects’ “increasing” responses to the graph t-value (Klein, 2001) and we found that this measure varied in the general population and reflected the participants’ self-evaluation of statistical knowledge (but not, crucially, their self-evaluation of first language skills). This suggests that the development of intuitive graphics skills correlates with math expertise, similarly to the positive effect of mathematical education on the accuracy of numerosity perception (Piazza et al., 2013). Whether a better grasp of numerical concepts strengthen graph-based statistical judgments or vice versa remains an open question that could be better addressed in the future through a finer assessment of participants’ numerical skills and their longitudinal follow-up. This is particularly necessary in the case of children: in our sample, we found a strong correlation between intuitive graphics skills and both arithmetic and reading performance, which does not allow to conclude for a specific correlation of intuitive graphics with numerical cognition. Interestingly, however, a relation between complex graph understanding and numerical cognition does indeed seem to exist (Ludewig et al., 2020). While a link between intuitive graphicacy and more complex statistical graph understanding remains to be shown, we believe that our trend judgment task could be an adequate assessment tool for the former, being simple and fast to perform. Similar to a previous numerosity perception test (Halberda et al., 2012), we have made the test publicly available online, so that it can be freely run by all researchers interested in investigating the correlation between their participants’ intuitive graphics skills and other abilities. Also, future research could determine if training on this task would generalize to higher-level graph understanding, in the same way that training intuitive number sense is suggested to increase arithmetic skills (Park & Brannon, 2013; but see Szkudlarek et al., 2021).

The second motivation of our study was to investigate whether intuitive graphics skills were available uniquely to individuals previously exposed to graphical representations, or whether they could be found in the absence of any such exposure. Children seem able to discriminate graphs based on the location and size of datapoints (Panavas et al., 2022), but no study to date tested whether they can perform statistical judgments on them. We show here that French 6-years-old children (unexposed to graphical representations) and uneducated Himba (living in an unindustrialized society in Northern Namibia where graphs are essentially absent) base their intuitive decisions on the *t*-value of the scatterplot. This shows that intuitive graphics skills are universally available and emerge early on in development, irrespectively of previous exposure to graphical representations. The present finding echoes previous data supporting the existence of a universal understanding of quantities (Dehaene, 2011; Pica et al., 2004), geometrical shapes (Sablé-Meyer et al., 2021), probabilities (Xu & Garcia, 2008), physics (Atran, 1998), and human psychology (Bjorklund, 2014). However, we found that the graphicacy index was much lower in Himba and children than in educated adults. Taken together, our results strongly suggest that intuitive graphics skills do not solely mature as a function of age, but can be refined by education and/or exposure to statistics and graphical representations. Recent evidence (Piazza et al., 2018) suggests that the precision of numerical estimations increases with education through an improved ability to focus on relevant information in the task (thus discarding non-numerical features). Future studies may investigate whether a progressive refinement of the filtering of irrelevant information (e.g., noisy or outlier datapoints) is also responsible for the relationship between the graphicacy index and education.

In summary, by investigating the premises of human intuitive graph perception, our study lays the foundations of a quantitative assessment of human graphicacy, an essential goal in building effective and early educational interventions that might in turn strengthen the comprehension of the complex graphs that humans are more and more routinely confronted with.

## ACKNOWLEDGMENTS

Funders: INSERM-CEA, Collège de France, Bettencourt-Schueller foundation, ERC grant “NeuroSyntax”, Mind Science Foundation Grant. We thank the 1^st^-grade children and teachers of Versailles’ academy; the Himba people and the local guides; all participants that took part in the online experiment; and Dario Mirossi, Mynie Zhan and Chanel Valera for translating the instructions of our online task.

## AUTHOR CONTRIBUTIONS

LC: conceptualization, data curation, formal analysis, funding acquisition, investigation, methodology, resources, software, visualization, writing (original draft), writing (review & editing) MSM: data curation, investigation, methodology, resources, software

EB: investigation

MJ: investigation

CPW: investigation

SC: data curation, investigation, project administration, writing (review and editing)

SD: conceptualization, formal analysis, funding acquisition, methodology, project administration, resources, software, supervision, validation, visualization, writing (review and editing)

## SUPPLEMENTARY MATERIALS

Correlations between the graphicacy index and other factors are briefly presented here. The graphicacy index correlated with participants’ self-evaluations of: their self-reported ability to understand scatterplots (r = .23, df = 3225, p < .0001); their familiarity with scatterplots (r = .23, df = 3217, p <.0001); and their mathematical knowledge (r = .22, df = 2028, p < .0001). The graphicacy index was also an inverted U shaped function of age: it increased until the age of ∼35 and then decreased. It also significantly increased with higher levels of formal education (r = .14, df = 3417, p < .0001). Concerning gender, no significant difference of the graphicacy index was observed between women and non-binary participants (women = 1.28, non-binary = 1.23, t(97.8) = 0.73, p = .47) but we found a significant, although small, advantage in favor of men over women (men = 1.54, t(2145.9) = 9.44, p < .0001) and over non-binary participants (t(109.8) = 4.28, p < .0001). Although these results are in agreement with previous research suggesting better spatial abilities in men than in women (for a review, Li et al., 2019), those results are inconclusive given that the present sample was self-recruited and not representative. Also, such difference was not found in the Himba (t(83.4) = 0.6, p = .55), nor in children (t(24.4) = 0.3, p = .77), thus suggesting that a higher ability to perform the task in online male participants could arise due to socio-cultural factors.

We found no significant difference in graphicacy index between participants responding on a touchscreen and on a computer (t(2510.9) = −.9, p = .37), nor between participants asked to respond with their left hand/finger for scatterplots going up and those asked to respond with their right hand/finger for scatterplots going up (t(3410.6) = .11, p = .91).

